# The effect of sequence learning on sensorimotor adaptation

**DOI:** 10.1101/2020.07.06.190397

**Authors:** Yang Liu, Hannah J. Block

## Abstract

Motor skill learning involves both sensorimotor adaptation (calibrating the response to task dynamics and kinematics), and sequence learning (executing the task elements in the correct order at the necessary speed). These processes typically occur together in natural behavior and share much in common, such as working memory demands, development, and possibly neural substrates. However, sensorimotor and sequence learning are usually studied in isolation in research settings, for example as force field adaptation or serial reaction time tasks (SRTT), respectively. It is therefore unclear whether having predictive sequence information during sensorimotor adaptation would facilitate performance, perhaps by improving sensorimotor planning, or if it would impair performance, perhaps by occupying neural resources needed for sensorimotor learning. Here we evaluated adaptation to a distance-dependent force field in two different SRTT contexts: In Experiment 1, 28 subjects reached between 4 targets in a sequenced or random order. In Experiment 2, 40 subjects reached to one target, but 3 force field directions were applied in a sequenced or random order. We did not observe any consistent influence of target position sequence on force field adaptation in Experiment 1. However, sequencing of force field directions facilitated sensorimotor adaptation and retention in Experiment 2. This is inconsistent with the idea that sensorimotor and sequence learning share neural resources in any mutually exclusive fashion. These findings indicate that under certain conditions, perhaps especially when the sequence is related to the sensorimotor perturbation itself as in Experiment 2, sequence learning may interact with sensorimotor learning in a facilitatory manner.

## 1 Introduction

In real-world motor skill acquisition, individuals must learn both sensorimotor and sequence aspects of the movement. Sensorimotor learning involves optimizing kinematic and dynamic parameters of a movement with sensory information. This involves learning aspects of the movement like forces and distances. For example, when a typist, fluent in typing on a modern small light-tough keyboard, is switched to an old-fashioned mechanical typewriter, typing speed and accuracy would likely drop due to sensorimotor factors such as wider spread of buttons and heavier resistances of keys. The process of compensating for such factors on a different keyboard is an example sensorimotor learning. In research, one of the classic sensorimotor learning tasks is force field adaptation (Shadmehr & Mussa-Ivaldi, 1994). The subject experiences externally-applied forces that will perturb a normal movement, such as reaching. Through trial and error practice, subjects learn to compensate for the forces, adapting to reduce reaching errors and recover their baseline level of performance. If the forces are abruptly removed, subjects who have adapted will display a negative aftereffect.

In contrast with sensorimotor learning, sequence learning involves learning a pattern of movements that are in a repeating order and optimizing the performance. Common examples could be learning to type words or phrases that come up frequently, such as one’s email address or name. This requires quickly producing the correct finger movements in the correct order. Classic laboratory sequence learning tasks such as serial reaction time task (SRTT) usually have minimal motor requirements, such as button pressing (Summers, 1975). The change in reaction time is a typical measure for SRTTs, with faster reaction time indicating sequence learning.

Sensorimotor learning and sequence learning are both critical for motor skill acquisition. In natural behavior, these processes most likely occur together. For example, learning the sequence of footsteps and the force of each step together in a dance routine. However, in the research setting, sequence and sensorimotor learning have largely been studied in isolation, limiting our understanding of how these processes interact. Sensorimotor and sequence learning have some properties in common, such as working memory demands and implicit vs. explicit components. Anguera, Reuter-Lorenz, Willingham, and Seidler (2010) found that spatial working memory is positively correlated with adaptation rate during the early stage of a visuomotor rotation task, illustrating the importance of working memory in sensorimtotor learning. Spatial working memory has also been shown to play a role in both explicit (Bo and Seidler 2009) and implicit (Bo, Jennett, and Seidler 2011) sequence learning.

Sensorimotor and sequence learning also rely on many of the same neural substrates. A meta-analysis comparing functional magnetic resonance imaging (fMRI) studies (Hardwick, Rottschy, Miall, & Eickhoff, 2013) found that both sequence and sensorimotor learning tasks activated left dorsal premotor cortex, left primary motor cortex (M1), supplementary motor area (SMA), and right cerebellum. In contrast, putamen was only active during sensorimotor learning, while thalamus was active during sequence learning alone. M1 is heavily involved in consolidation and retention periods of force field adaptation (Richardson et al., 2006; Hunter, Sacco, Nitsche, & Turner, 2009; Galea, Vazquez, Pasricha, Orban de Xivry, & Celnik, 2011). Premotor cortex is engaged in sequence learning (1998; 2000) as well as sensorimotor learning (Kurata & Hoshi, 1999). SMA plays an important role in programming complex sequential movements (Roland, Larsen, Lassen, & Skinhoj, 1980) and its activity increases throughout the learning process (Grafton, Hazeltine, & Ivry, 1995; Toni et al., 1998). SMA proper is also engaged in sensorimotor learning tasks (Gerlo & Andres, 2002; Serrien, Strens, Oliviero, & Brown, 2002). Previous studies also found cerebellum involvement in both adaptation (Jayaram et al., 2012; Galea et al., 2011) and sequence learning (2002) tasks.

While much can be deduced by comparing the results of sequence learning and sensorimotor learning studies, many questions can only be answered by developing tasks in which sensorimotor and sequence learning can be studied simultaneously, in the same participants. For example, knowing that both processes depend on working memory and certain brain regions does not tell us whether they interfere with or facilitate each other when occurring at the same time. It is unknown whether having predictive sequence information during sensorimotor learning facilitates performance, perhaps by improving sensorimotor planning, or if it impairs performance, perhaps by occupying neural resources needed for sensorimotor learning.

Here we asked whether the addition of a sequence component facilitates sensorimotor learning in healthy adults. If sensorimotor learning improves when there is an embedded sequence component, it would suggest a beneficial interaction that could one day be taken advantage of in motor rehabilitation for clinical populations with impaired movement, such as stroke. If sensorimotor learning worsens when there is a sequence component, it would suggest a detrimental interaction, perhaps due to the monopolizing of shared resources, that requires further study. Should sequence learning have no effect on sensorimotor learning, it would suggest these two motor skill systems operate independently in behavior. Because it is unknown what type of sequence component might affect sensorimotor learning, we tested two. Experiment 1 tested the effect of sequenced target positions in force field adaptation, while Experiment 2 tested the effect of sequenced force field directions on force field adaptation.

## 2 Materials and Methods

### 2.1 Participants

Typically developing adults participated in both experiments. Twenty-eight individuals aged 24.11 ± 6.22 years old participated in Experiment 1, and 40 adults aged 22.50 ± 4.72 years old participated in Experiment 2 (mean ± SD). All participants reported no history of neurological disorders or upper limb muscular injuries. All participants gave written informed consent. The study was approved by the institutional review board of Indiana University Bloomington.

### 2.2 Apparatus

Participants performed reaching movements on a robotic apparatus (KINARM End-Point Lab, BKIN Technologies, Kingston, Canada). The KINARM system uses a 2D virtual reality display to present visual stimuli (Figure 1.A). A downward facing TV on top projected tasks onto the mirror. Participants were seated in front of the apparatus and grasped the robotic manipulandum. Participants viewed the task display in the mirror, which prevented vision of the hand and manipulandum. The task display appeared to be in the same horizontal plane as the robotic manipulandum. A drape over the shoulders prevented participants from seeing their upper arm and shoulder. Veridical visual feedback of the hand’s position was provided as a white dot in the task display. For experiment 1, electromyography (EMG) was collected using an 8-channel EMG system from Bortec (AMT-8). The reaching tasks were programmed using Simulink toolbox from MATLAB 2017b (MathWorks Inc., Natick, MA, United States). Tasks were operated, and data was stored through the operation and acquisition software Dexterit-E by BKIN Technologies. The sampling rate for hand position and force was 2,000HZ.

**Figure 1:**
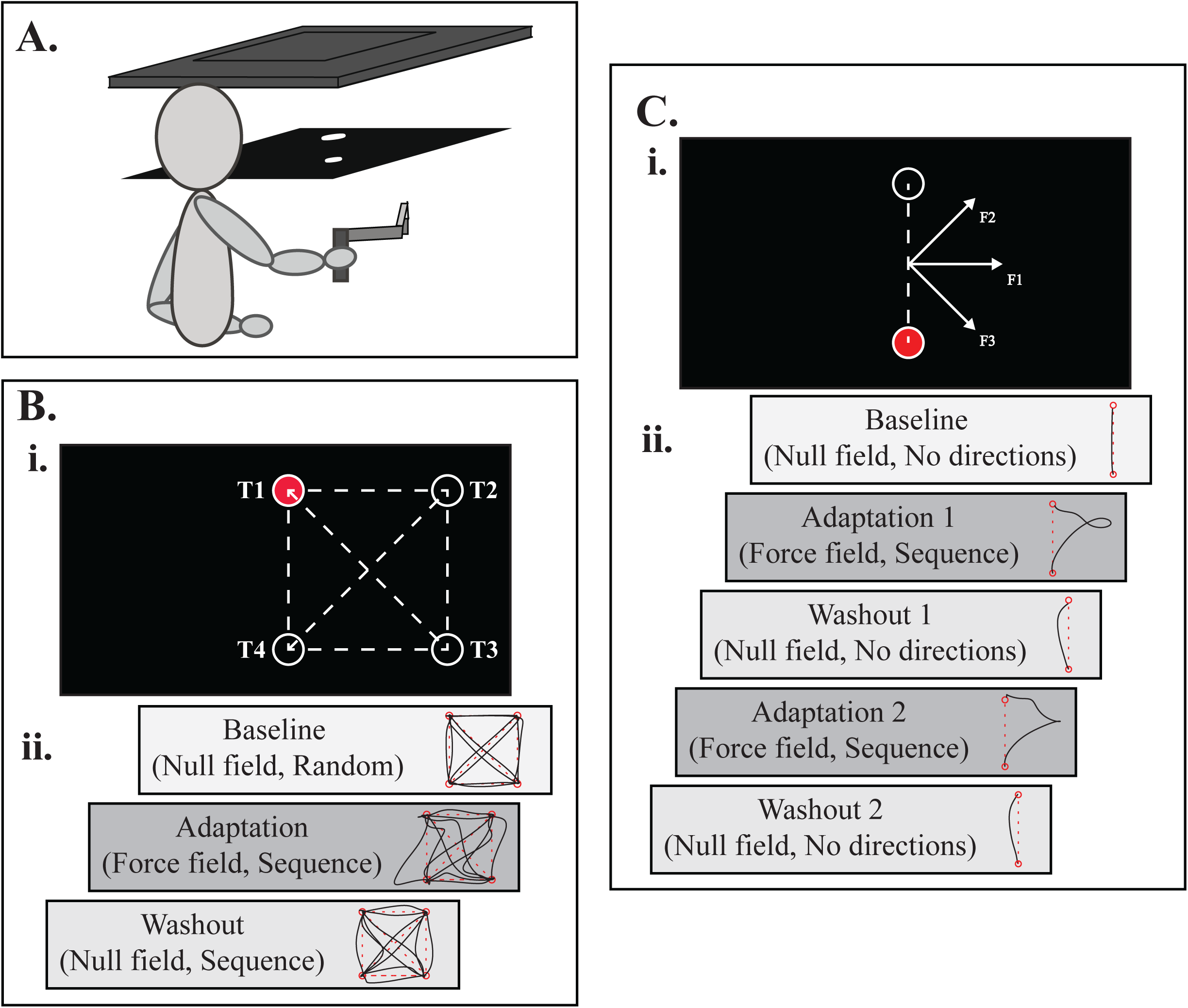
A. KINARM End-Point apparatus setup. Participants grasped the manipulandum and viewed tasks from the mirror. B. Experiment 1 study design. i. Task layout for a right-handed participant. T1-4: labels for the four target positions. Red target was the start position for each trial. The 12-element sequence was: T1-T4 (Reach 1), T4-T2 (Reach 2), T2-T3 (Reach 3), T3-T4 (Reach 4), T4-T1 (Reach 5), T1-T3 (Reach 6), T3-T1 (Reach 7), T1-T2 (Reach 8), T2-T4 (Reach 9), T4-T3 (Reach 10), T3-T2 (Reach 11), T2-T1 (Reach 12). ii. Task blocks for sequence targets with force field group and example movement paths of a participant’s first trial in each block. C. Experiment 2 study design. i. Task layout for a right-handed participant. Red target was the start position for each reach. F1 was the perpendicular force direction; F2 was the 45° force direction; F3 was the 315° force direction. The 6-element sequence was: F1-F2-F3-F1-F3-F2. ii. Task blocks for sequence directions with force field group and example movement paths of a participant’s first reach in each block.

### 2.3 Force field design

Force field adaptation in reaching is commonly studied with a velocity dependent force field perturbation, where the amount of force is directly related to the subject’s movement velocity (Shadmehr & Mussa-Ivaldi, 1994). However, this type of force field could create a confound in a sequence-learning study: Subjects might move faster as they learn the sequence, which would result in stronger force perturbations, interfering with sensorimotor learning. We therefore developed a novel distance-dependent force field, designed to apply the same pattern of forces regardless of subjects’ movement velocity.

First we computed the vector from the end target to start target (SE):

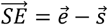

where e and s were the vectors for start and end targets; we computed γ, which was the force angle perpendicular to the SE vector:

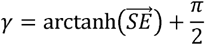

*f*x and *f*y were the lateral (x) and sagittal (y) directions of the maximum manipulandum force magnitudes:

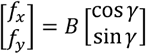

where *B* is the force viscosity, usually in unit *N*; cos θ was the angle between the reaching hand and the start target and the start and end targets.

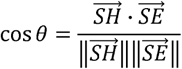

where SH was the vector for start target and reaching hand positions. The ratio of the projection of the reaching hand onto the line defined by the start and end targets was then determined. *f’*x and *f’*y were the manipulandum forces at hand position:

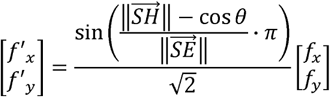

To mimic the reaching pattern of a velocity-dependent force field, we scaled the force field into a bell shape. The force field was gradually introduced as the reaching hand left the start target, maximized half-way through the reach, and gradually dialed down as the hand got closer to the end target. Meanwhile, in spite of individual reaching speed differences, the same amount of external perturbation was applied to each participant with the same ratio between the start and end targets. In other words, participants experienced the same scale of external force during reaching regardless of hand movement velocity or force direction. Participants did not experience any force or resistance while their hand was in a target.

### 2.4 Experimental Procedure

After signing the informed consent form, participants completed the Edinburgh Handedness Inventory (Oldfield, 1971) to determine which hand would be the dominant hand in performing the reaching task. Participants were randomly assigned to one of the groups in the experiment. If the participant was assigned to experiment 1, EMG surface electrodes were placed on seven muscles of the participant’s dominant arm. The muscles were: brachioradialis, anterior deltoid, medial deltoid, posterior deltoid, biceps long head, triceps long head, and pectoralis clavicular.

Participants were seated in front of the robotic apparatus (Figure 1.A). They were instructed to move the robotic manipulandum briskly to a target in a straight line. When a target turned green, it signaled the participant to start reaching towards it. Once the hand arrived at the target, the color turned red and participants were asked to hold the manipulandum within the target until the next target turned green. A real-time white dot representing the participant’s hand was visible during both experiments. Participants were familiarized with the task by a practice block with no force or sequence.

Participants were not informed of when and where the force field would be onset and participants were also not given any hint regarding the possible existence of a sequence. The experimenter asked participants whether they noticed any pattern or sequence upon experiment completion. If answered yes, the experimenter would ask the participants to describe what they noticed. For experiment 1, if the participant noticed sequence of reach directions or target appearances, the experimenter would provide an image of the task layout (Figure 1.B.i) and ask the participant to draw the remembered sequence. For experiment 2, if participants noticed the different force directions or sequence of the force directions, the experimenter recorded the order they recalled.

#### 2.4.1 Experiment1

The task displayed four 1cm-diameter hollow circles as the targets on the screen, located at the four corners of a 20cm square (Figure 1.B.i). Two targets were displayed on the participant’s midline and the other two targets were on the dominant hand-side of the display. In total, the reaching task had 40 trials, each trial consisting of 12 reaching movements. Thus, the entire task contained 480 reaching movements. The 12 different directions of the square -- vertical, diagonal, and horizontal -- cover all the reaching directions of a trial (Figure 1.B.i). Each trial started and ended at the same target (T1, top mid-line target). The sequence element was implemented in the target order. Upon sequence onset in the adaptation block, the targets in each reaching trial appeared in a fixed order. After each trial, the total movement time was displayed on the screen to give participants a general idea about their speed. This served as a reminder to avoid significant slowing down later in the experiment due to fatigue or boredom.

Participants were randomly assigned to a sequence with force group (SF, n = 16) where the set of targets appeared in a particular order and participants experienced a force field, or a random with force group (RF, n = 12) where targets appeared in a random order, but participants still experienced a force field. The first block was a baseline consisting of five trials in a null field, with the targets presented in random order. The second block was an adaptation block (25 trials). During the adaptation block, targets appeared in a fixed order in the SF group, whereas targets appeared randomly in the RF group. Both the SF and RF groups experienced force perturbations during the adaptation block. A catch reach was implemented in this block to confirm subjects were attending: the external force was turned off without notification to the participants to detect the aftereffects in this reach (Shadmehr & Brasher-Krug, 1997). The washout block contained 10 trials with no force perturbation; the SF group continued to experience the same sequence (Figure 1.B.ii).

#### 2.4.2 Experiment 2

All reaches were straight-ahead, with two targets aligned on the participant’s midline. The start target was closer to the participant and the end target was 20cm forward of the start target. Participants were instructed to make straight and brisk movements from start to end target. After reaching the end target, the robot automatically brought the testing arm back to the start target for the next trial. There were three force directions pushing to the dominant hand side of the participant: a horizontal direction (0° perpendicular to the straight line between start and end targets), a 45° upward direction, and a 315° downward direction (Figure 1.C.i).

Participants were randomly assigned to one of two groups: a sequenced force directions group (SDF) and a random force directions group (RDF). The task was divided into five blocks. The first was a baseline block where no force was applied (24 reaches). The baseline was followed by an adaptation block (180 reaches), in which both groups experienced a distance-dependent force field. Every 6 reaches in the two adaptation blocks comprised the three force directions, each repeating twice. For the SDF group, the three directions of force were in a repeating order of 6 elements. Adaptation was followed by the first washout block (42 reaches), in which no forces were applied (null force). In the second adaptation block (90 reaches), the settings of force field and directions were the same as the first adaptation block. The second washout block (24 reaches) was performed in the null field (Figure 1.C.ii). For 17 participants, the baseline block was 20 reaches, washout 1 was 40 reaches, and washout 2 was 20 reaches.

### 2.5 Outcome variables and analysis

All data were preprocessed and analyzed using Matlab. Outcome measures from the manipulandum data were maximum perpendicular deviation and reaction time. Perpendicular deviation is the distance between the hand reaching trajectory to the straight line between the two targets. The maximum perpendicular deviation can thus be used as a measure of sensorimotor error (Darainy & Ostry, 2008). Perpendicular deviation was averaged every 12 reaches (one sequence) in experiment 1 and every 6 reaches (one sequence) in experiment 2. Perpendicular deviations in adaptation and washout blocks were normalized by subtracting the last trial/reach in the baseline block. Reaction time was measured as the time from target onset to movement being detected (the moment the hand left the start target). To compare the rate of adaptation during force field blocks, we defined early adaptation to be reaches 2 - 11 in the adaptation blocks in both experiments (Lamothe et al., 2014). We did not include the first reach because the unexpected onset of the force field could surprise subjects. We calculated the linear slope of reaches 2-11 as the early adaptation rate (Diedrichsen, 2007).

EMG was measured as another way to assess sequence learning in Experiment 1. Preprocessing techniques including de-trending, rectifying, and low frequency band pass filtering were adopted to remove baseline drift in EMG data. Anticipatory EMG was defined by the time window between 100ms prior to the movement onset and the moment of movement onset and was integrated (IEMG) (Lang & Bastian, 1999). Anticipatory IEMG is the amount of muscle activity the participant used to hold the robotic manipulandum in the start position. Normalization was done by dividing anticipatory IEMG by the IEMG during baseline.

Two-way mixed model analysis of variance (ANOVA) with repeated measures was used to analyze change of reaction time from pre to post sequence learning. The between-subjects factor “group” was the SF and RF groups in experiment 1 and the SDF and RDF groups in experiment 2. The within-subjects factor “time” was the first and last trials in the adaptation blocks. To compare max perpendicular deviation, two-way mixed model ANOVA with repeated measures was performed on adaptation and washout blocks, with within-subjects factor “time” to be the trials in block and between-subjects factor “group” to be the groups in each experiment. Post-hoc pairwise comparison using Tukey honest significant difference test was performed if significance was found after performing ANOVA. For experiment 1 we extracted each muscle’s normalized anticipatory IEMG for each reach direction and performed the same repeated-measures ANOVA to compare across groups for each muscle. We did not match the IEMG in the RF group to the SF group in reaching directions as the participants in the RF did not have the sequence information to predict the upcoming reaching directions. The trial that had an unexpected catch reach was not included in the EMG analysis.

## 3 Results

### 3.1 Experiment 1

#### 3.1.1 Sensorimtor learning

Figure 1.B.ii shows movement paths of an example participant. During the baseline block, the movement paths showed very few errors. When the force field was first onset in the adaptation block, the movement paths deviated from the straight line between targets; after trial-and-error practice, movement paths in the last trial of the adaptation block showed fewer errors, similar to the path pattern in the baseline trial, which indicates adaptation occurred. When the external force was just turned off at the beginning of the washout block, we observed large deviations to the opposite side of the first adaptation trial, reflecting a negative aftereffect.

Comparing normalized perpendicular deviation per trial (error averaged across reach direction) between the SF and RF groups revealed no differences in sensorimotor learning (Figure 2.A). In the adaptation block there was a significant main effect of time (*F*(24, 624) = 63.51, *p* < 0.0001), consistent with trial-by-trial error reduction. However, there was no effect of group (*F*(1, 624) = 0.41, *p* = 0.53) or group by time interaction (*F*(24, 624) = 0.85, *p* = 0.68). The washout block also had a main effect of time (*F*(9,234) = 180.05, *p* < 0.0001), consistent with a negative aftereffect. There was no group (*F*(1, 234) = 0.37, *p* = 0.55) or group by time interaction (*F*(9, 234) = 0.34, *p* = 0.96) for the washout block. Comparing the catch reach between the two groups found no significant difference (*t*(26) = -1.51, *p* = 0.14). Early adaptation rate was not significantly different between the two groups either (*t*(26) = -0.45, *p* = 0.66) (Figure 2.B).

**Figure 2:**
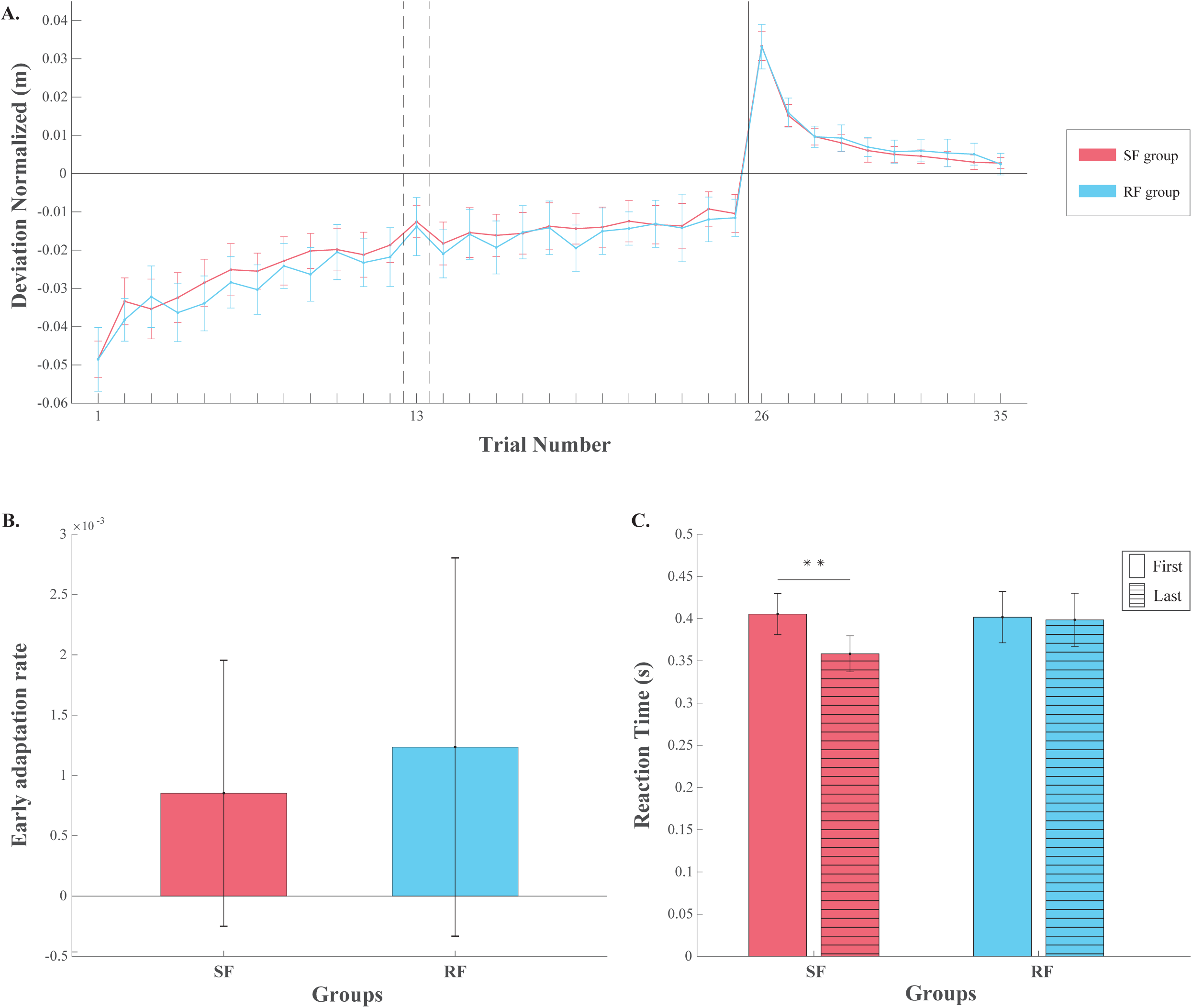
Experiment 1. A. Normalized group mean sensorimotor errors across trials. Perpendicular error in each trial, averaged across directions and normalized to baseline. Trial number 1-25: adaptation block. Trial number 26-35: washout block. Dotted line indicated the trial that had the catch reach. B. Early adaptation rate. C. Reaction time (RT) in the first trial of adaptation and last trial of washout. * Significant time x group interaction (p < 0.01), suggesting RT decreased in the SF group relative to the RF group. Error bars: 95% confidence interval.

Examining individual reach directions, rather than averaging perpendicular error across the 12 directions in each trial, revealed some group differences (Figure 3). Among the four vertical reach directions (Figure 3.A), there was a significant interaction of group and time for reach direction 5 in washout (*F*(9, 234) = 2.05, *p* = 0.035). There was also a significant main effect of group for reach direction 1 in washout (*F*(1, 234) = 8.22, *p* = 0.0081). Significant main effects of time were observed in adaptation (*p* < 0.0001) as well as washout (*p* < 0.0001). Among the four diagonal reach directions (Figure 3.B), a significant group and time interaction was found for reach direction 9 in adaptation (*F*(24, 624) = 1.83, *p* = 0.0094). For reach directions 2, 7, and 9, the main effects of group were significant in adaptation (*F*(1, 624) = 4.59, *p* = 0.042; *F*(1, 624) = 8.98, *p* = 0.0059; *F*(1, 624) = 4.27, *p* = 0.049). The main effects of group for reach directions 2 and 7 were also significant in washout (*F*(1, 234) = 4.20, *p* = 0.050; *F*(1, 234) = 4.53, *p* = 0.043). All four diagonal reach directions had significant main effects of time in both adaptation (*p* < 0.0001) and washout (*p* < 0.0001). Among the four horizontal reach directions (Figure 3.C), a significant interaction of group and time was found for reach direction 8 in adaptation (*F*(24, 624) = 1.65, *p* = 0.027). For reach directions 4, 8, and 10 in adaptation, there were significant main effects of time (*F*(24, 624) = 1.77, *p* = 0.014; *F*(24, 624) = 1.88, *p* = 0.0071; *F*(24, 624) = 8.03, *p* < 0.0001). All four horizontal reach directions had significant main effects of time in washout (*p* < 0.0001). All reaches were corresponding to the reaching order in the sequence group (Figure 1.B).

**Figure 3:**
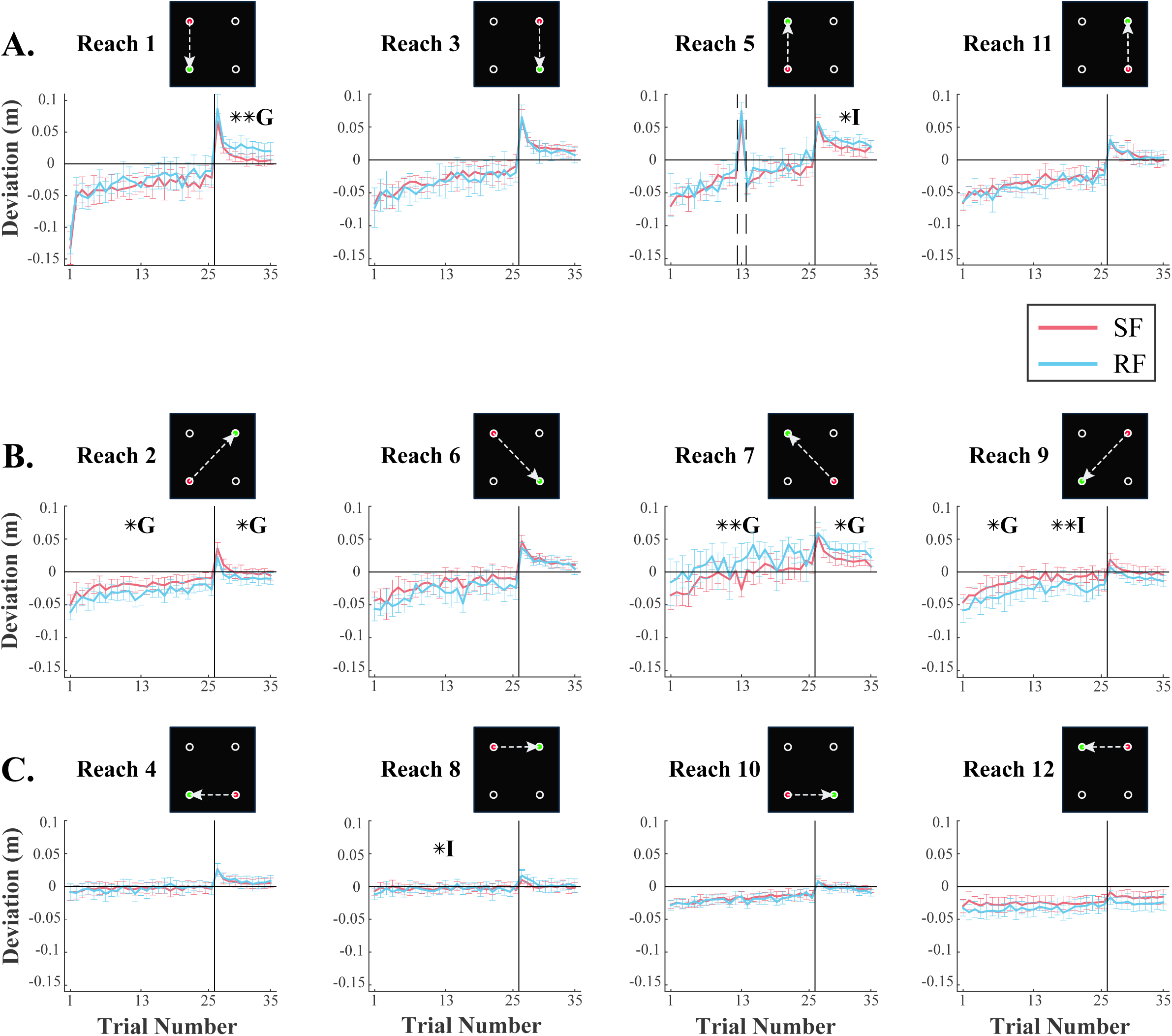
Normalized group adaptation curves for 12 individual reach directions. Trial number 1-25: adaptation block. Trial number 26-35: washout block. A: Vertical reaches. Dotted line in reach direction 5 indicated the catch reach. B: Diagonal reaches. C: Horizontal reaches. G: statistically significant main effect of group in the adaptation or washout blocks. I: statistically significant interaction of group and time in the adaptation or washout blocks. Error bars: 95% confidence interval. * p < 0.05. ** p < 0.01.

#### 3.1.2 Sequence learning

13 out of 16 participants in the SF group were able to recognize at least part of the sequence, the average number of sequence elements recognized was 5.3125 ± 2.5348 (mean ± CI) in the SF group. Reaction time of first (onset of sequence exposure) and last (final repetition of sequence) trial in sequence learning were compared between the SF and RF groups. There was a significant interaction of time and group (*F*(1, 26) = 8.29, *p* = 0.0079), suggesting the SF group’s reaction time decreased due to sequence learning relative to the RF group (Figure 2.C). There was a main effect of time (*F*(1, 26) = 13.98, *p* = 0.00092), but no effect of group (*F*(1, 26) = 1.38, *p* = 0.25).

The anticipatory IEMG results also supported the presence of sequence learning in the SF group compared to the RF group. This is indicated by group and time interactions in adaptation or washout blocks for at least one reach direction in most of the muscles we recorded from. For the brachioradialis muscle, there was a significant interaction of group and time in reach direction 4 during adaptation (*F*(23,598) = 1.58, *p* = 0.043). For the anterior deltoid muscle, there were significant interactions of time and group in reach directions 2-5, 7, 9, and 10 in the adaptation block (*p* < 0.05), and reach directions 1-6, 8, 10, and 12 in the washout block (*p* < 0.05) (Figure 4.ii). The middle deltoid muscle had significant interactions of time and group for reach directions 1, 2, and 5 in the adaptation block (*p* < 0.05) (Figure 4.iii), and reach directions 4, 6, and 8-10 in the washout block (*p* < 0.05). The posterior deltoid muscle had a significant interaction of time and group for reach direction 5 in the adaptation block (*F*(23,598) = 2.02, *p* = 0.0034) (Figure 4.iv) and reach direction 7 in the washout block (*F*(9,234) = 2.07, *p* = 0.033). For the triceps long head, there were significant interactions for reach directions 2-5, 7, and 9-11 in the adaptation block (*p* < 0.05), and reach directions 4-6, 8, 10, and 12 in the washout block (*p* < 0.05) (Figure 4.vi). The pectoralis clavicular muscle had significant interactions of time and group in reach directions 6 and 12 in the washout block (*p <* 0.05). The biceps long head had no significant interaction between group and time in anticipatory IEMG for any of the 12 reach directions in either the adaptation or washout blocks.

**Figure 4:**
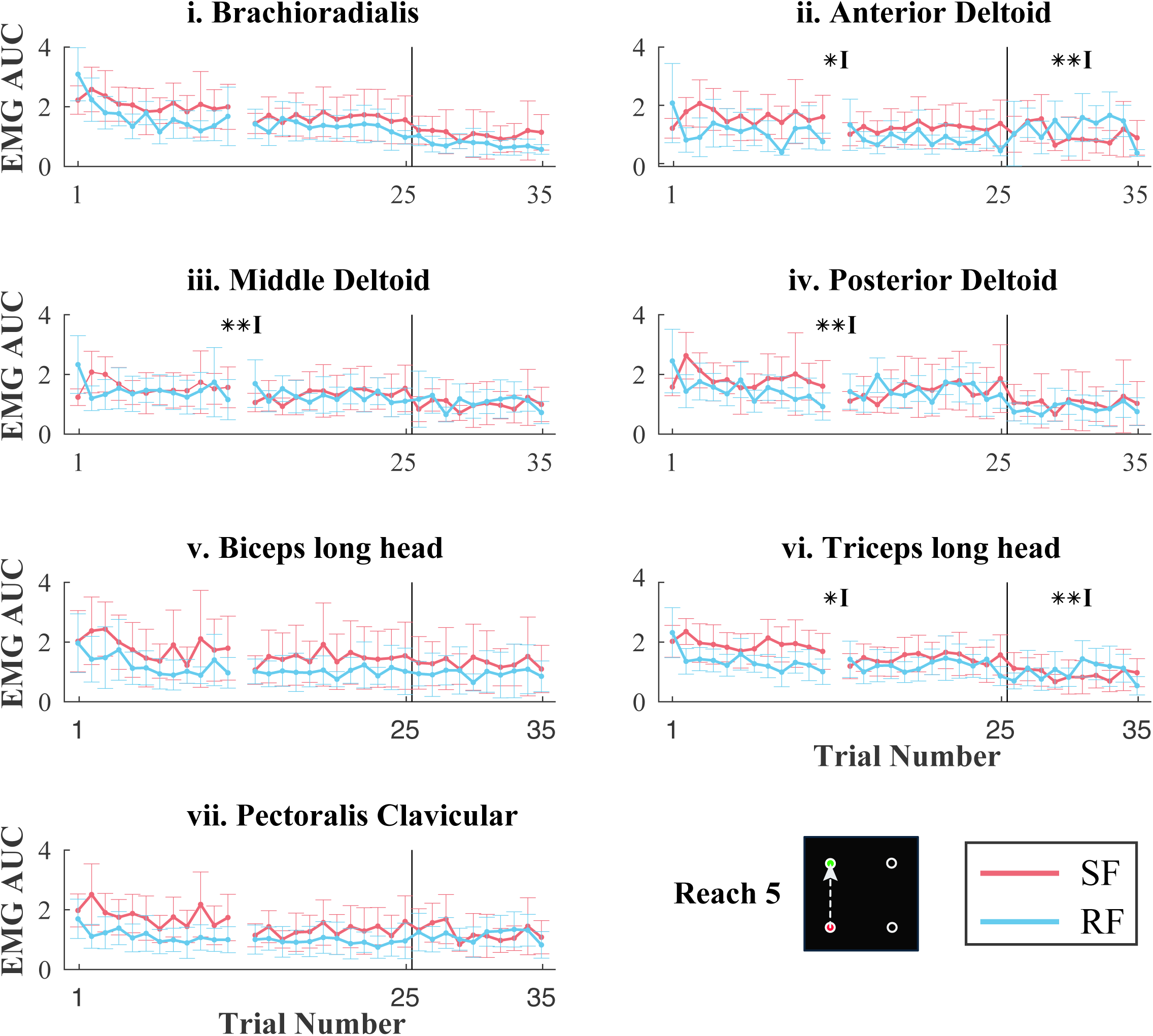
Example normalized anticipatory IEMG of 7 muscles: reach direction 5. Trial 1-25: adaptation block (All reaches in trial 13 were excluded due to the catch reach). Trial 26-35: washout block. I: statistically significant interaction of group and time in the adaptation or washout blocks. Error bars: 95% confidence interval. * p < 0.05. ** p < 0.01.

### 3.1 Experiment 2

#### 3.2.1 Sensorimotor learning

Figure 1.C.ii shows example movement paths of a participant in each block of Experiment 2. Little movement error was observed during the baseline block. When the force field was first onset in the two adaptation blocks, hand trajectories were largely deviated from the straight line; through trial-and-error practice, the last three trials in the adaptation showed smaller deviation from the straight line. When the external force was just removed in the two washout blocks, large deviations were observed to the opposite side of the adaptation trials, indicating a negative aftereffect.

We compared perpendicular deviation between the two groups in the baseline block and found no effect of time (*F*(3,114) = 2.22, *p* = 0.089), group (*F*(1,114) = 0.06, *p* = 0.81) or group and time interaction (*F*(3,114) = 0.98, *p* = 0.40). We then normalized each trial in the remaining four blocks by subtracting individuals’ mean baseline perpendicular deviation. Group adaptation curves are shown in figure 5.A. There was a main effect of time in the first adaptation block (*F*(29,1102) = 65.39, *p* < 0.0001), reflecting that both groups reduced perpendicular error across trials as expected. A significant interaction between group and time in the first adaptation block (*F*(29,1102) = 1.64, *p* = 0.018) suggests that the two groups reduced error at different rates. There was no main effect of group (*F*(1,1102) = 0.38, *p* = 0.54). In the second adaptation block, there was a significant main effect of time (*F*(14,532) = 57.29, *p* < 0.0001), indicating further adaptation took place, but no group effect (*F*(1,532) = 0.32, *p* = 0.57) or interaction (*F*(14,532) = 1.00, *p* = 0.45).

**Figure 5:**
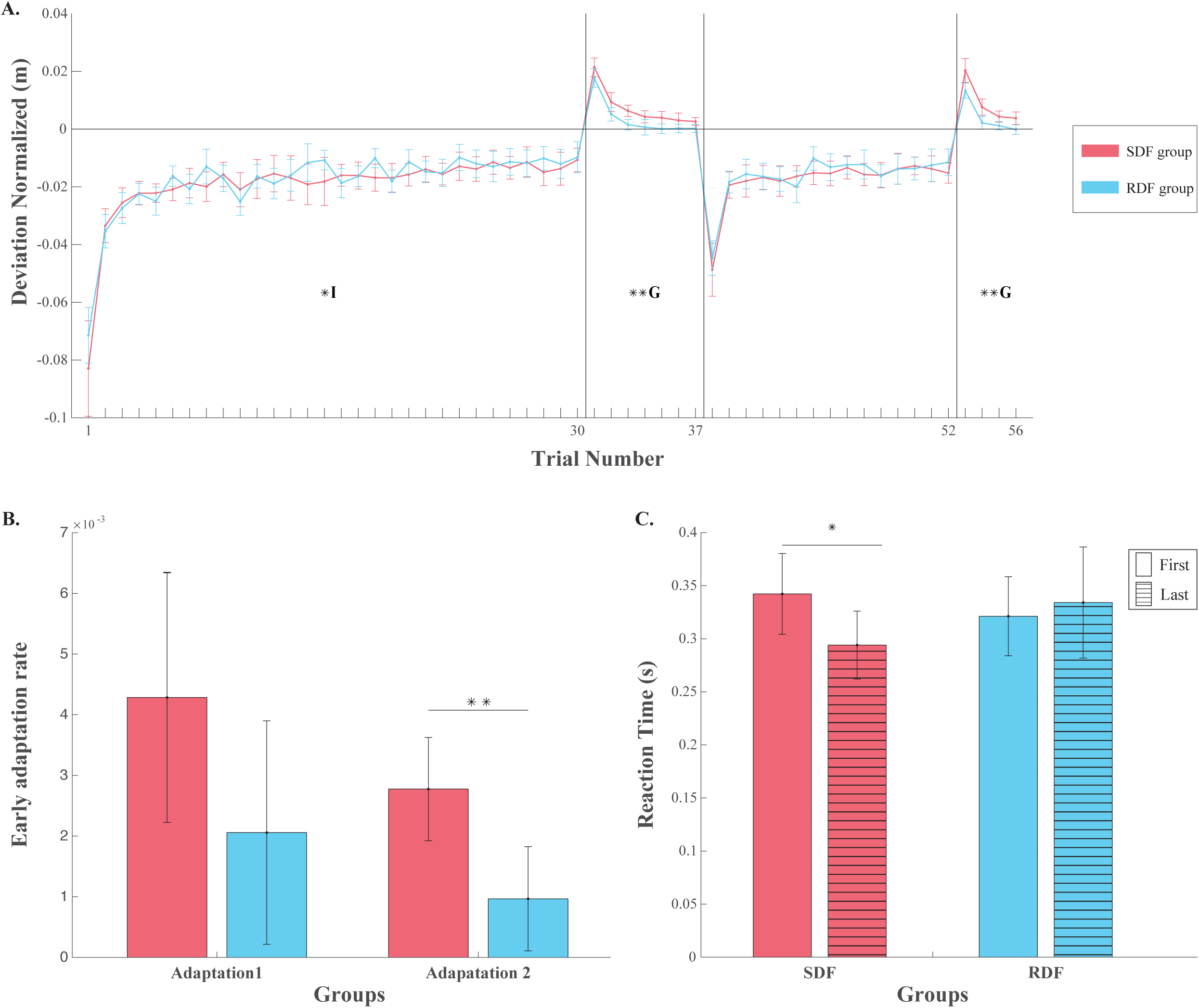
Experiment 2. A. Group mean sensorimotor errors across trials. Maximum perpendicular deviation was normalized to baseline. Trial number 1-30: adaptation 1. Trial number 31-37: washout 1. Trial number 38-52: adaptation 2. Trial number 53-56: washout 2. G: statistically significant main effect of group in the adaptation or washout blocks. I: statistically significant interaction of group and time in the adaptation or washout blocks. B. Early adaptation rate. ** p < 0.01. C. Reaction time in the first and last trial of sequence learning. * Significant time x group interaction (p < 0.05), suggesting RT decreased in the SDF relative to RDF group. Error bars: 95% confidence interval.

In the first washout block, there were main effects of time (*F*(6,228) = 147.58, *p* < 0.0001), consistent with a negative aftereffect, and group (*F*(1,228) = 8.76, *p* = 0.0053), with the sequence group appearing to have a more enduring negative aftereffect. There was no interaction (*F*(6,228) = 0.58, *p* = 0.75). In the second washout block, there was again a main effect of time (*F*(3,114) = 132.69, *p* < 0.0001) and group (*F*(1,114) = 13.23, *p* = 0.00081), but no interaction (*F*(3,114) = 1.99, *p* = 0.12).

Early adaptation rate was not significant between sequence and random groups in the first adaptation block (*t*(38) = 1.69, *p* = 0.10), but early adaptation rate was found to be significant in the second adaptation block (*t*(38) = 3.13, *p* = 0.0034). The early adaptation rate between the two adaptation blocks in either SDF or RDF groups was not significant (*t*(19) = 1.64, *p* = 0.12; *t*(19) = 1.28, *p* = 0.22) (Figure 5.B).

We also separately examined the perpendicular deviation from each of the three force directions in the two adaptation blocks (Figure 6). In the 0° horizontal force direction reaches (F1, Figure 1.C.i), we found significant main effect of time for both adaptation blocks (*F*(19,2242) = 27.62, *p* < 0.0001; *F*(19,1102) = 12.38, *p* < 0.0001), suggesting subjects in both groups adapted to this force field direction. We also found main effects of group (*F*(1,2242) = 11.98, *p* = 0.00075; *F*(1,1102) = 9.71, *p* = 0.0028), and group by time interactions (*F*(19,2242) = 2.58, *p* = 0.00020; *F*(19,1102) = 2.49, *p* = 0.00039), suggesting differences between the sequence and random groups.

**Figure 6:**
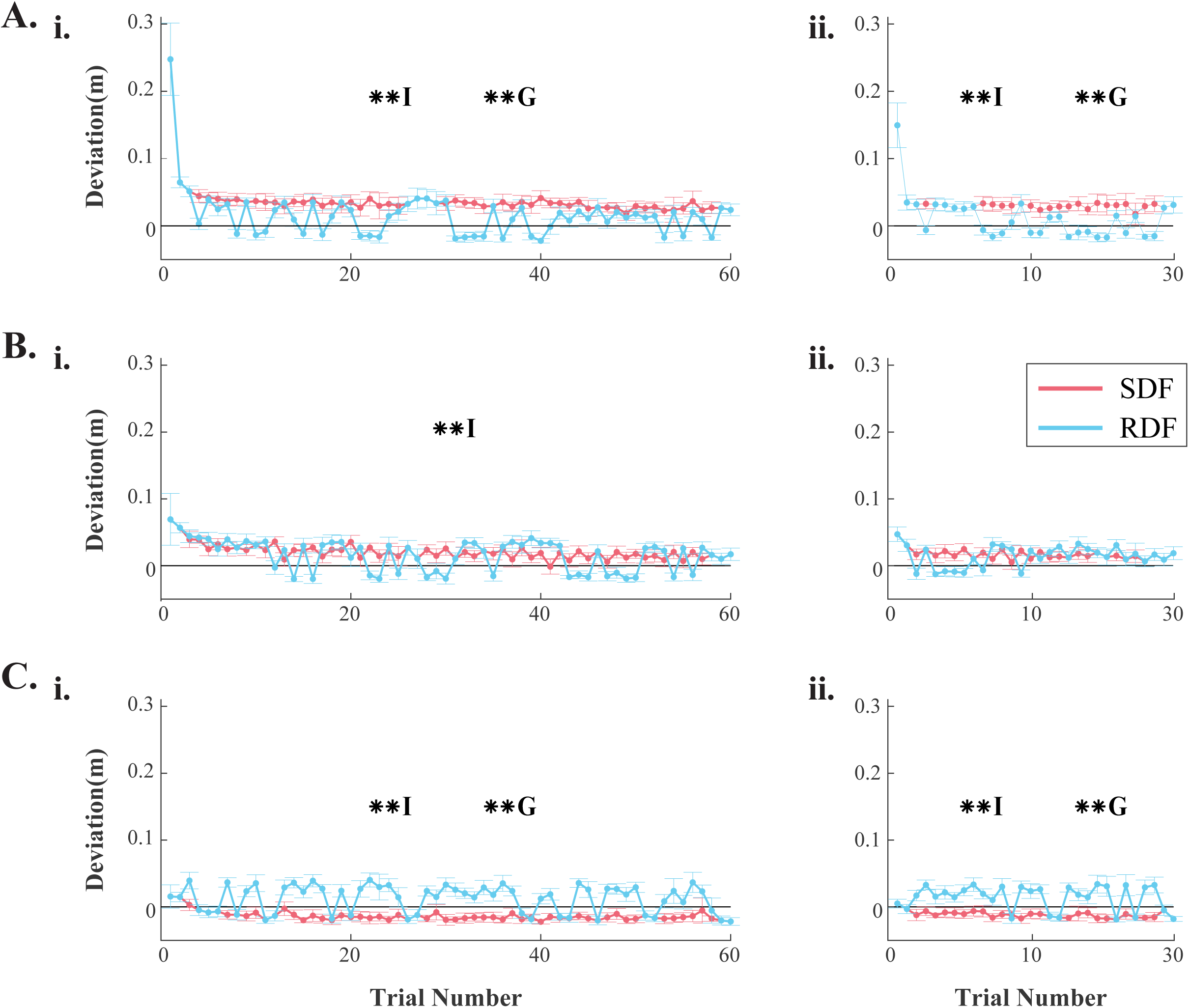
Adaptation curve of each force directions. A: 0° force direction. B: 45° force direction. C: 335° force direction. i. Adaptation 1. ii. Adaptation 2. G: statistically significant main effect of group in the adaptation blocks. I: statistically significant interaction of group and time in the adaptation blocks. Error bars: 95% confidence interval. **p < 0.01.

In the 45° upright force direction reaches (F2, Figure 1.C.i), both adaptation blocks had a main effect of time (*F*(19,2242) = 25.82, *p* < 0.0001; *F*(19,1102) = 17.41, *p* < 0.0001), but no effect of group (*F*(1,2242) = 2.16, *p* = 0.14; *F*(1,1102) = 0.85, *p* = 0.36). There was a group by time interaction in the first adaptation block (*F*(19,2242) = 1.92, *p* = 0.0094), but not the second adaptation block (*F*(19,1102) = 0.72, *p* = 0.80).

In the 335° downright force direction reaches (F3, Figure 1.C.i), we found significant interactions between group and time (*F*(19,2242) = 4.17, *p* < 0.0001; *F*(19,1102) = 3.77, *p* < 0.0001), main effects of time (*F*(19,2242) = 11.55, *p* < 0.0001; *F*(19,1102) = 9.41, *p* < 0.0001), and main effects of group (*F*(1,2242) = 86.49, *p* < 0.0001; *F*(1,1102) = 52.43, *p* < 0.0001) in both adaptation blocks.

#### 3.2.2 Sequence learning

In both groups, many participants noticed the forces were different, but only a few perceived that there were three force directions. 17 out 20 participants in the SDF group noticed different force directions, the average number of directions recognized in the SDF group was 1.90 ± 0.48 (mean ± CI). 7 SDF participants were also able to recall at least half of the sequence. In the RDF group, 18 out 20 participants noticed the different force directions, on average, participants in the RDF group were able to notice 1.55 ± 0.39 different force directions. We compared reaction time across SDF and RDF groups between the first trial (6-element sequence onset) in adaptation 1 and the last trial in adaptation 2 (sequence ended). A significant interaction of time and group was found (*F*(1, 38) = 4.88, *p* = 0.033), suggesting reduced reaction time (sequence learning) in the SDF group relative to the RDF group (Figure 5.C). There was no main effect of time (*F*(1, 38) = 1.63, *p* = 0.21) or group (*F*(1, 38) = 0.16, *p* = 0.69).

## 4 Discussion

Here we asked whether adding a sequence component would facilitate sensorimotor learning. We tested this idea in two ways: Sequencing of target positions in space (experiment 1) and sequencing of force field directions (experiment 2). We did not observe any consistent influence of target position sequence on force field adaptation in experiment 1. However, sequencing of force field directions facilitated the adaptation speed and increased retention in experiment 2. These findings indicate that under certain conditions, sequence learning may interact with sensorimotor learning in a facilitatory manner.

### 4.1 Reaction time decreased among sequence groups during reaching

In the current study, we observed decrease in reaction time for the sequence, but not random, group in both experiments. This suggests that subjects were able to learn the sequence, whether it was a sequence of target positions in experiment 1 or of force field directions in experiment 2. Classic button-press serial reaction time tasks (Nissen & Bullemer, 1987; Willingham, Nissen, & Bullemer, 1989) evaluate sequence learning as the change in response time, which is movement onset time (OT) plus movement time (MT). However, sequence learning is difficult to distinguish from skill improvement in such a paradigm, as the change in response time is the result of both OT and MT. More recent approaches are able to distinguish sequence learning from sensorimotor skill learning by making the time of movement onset and the end of the movement into two recordable events. This allows examination of OT for sequence learning and MT for sensorimotor learning. For example, Moisello et al. (2009) conducted a sequence learning study using visuomotor reaching. They successfully tracked both OT and MT by having subjects move their finger from one target to another on a tablet, with the targets appearing either sequentially or randomly. Though they found a similar pattern of changes in response time to classic button-press SRTTs, the decrease in response time was largely due to skill improvement, as the MT reduced during sequence learning. Meanwhile, the increase of response time in a random trial during sequence learning was due to the increase of OT. This means response time might not be a good measure of sequence learning in reaching tasks.

In the current study, we used a robotic reaching tasks to extract OT (reaction time) out of response time, so we were able to investigate the sequence learning component in the force field adaptation task. In experiment 1, we used a 12-element sequence of target positions in space. While the sequence was relatively long, it was not directly related to the sensorimotor learning aspect of the task (i.e., force field adaptation). In addition, the presence of four target positions, if not the exact sequence of their presentation, was clear to participants. Therefore, in experiment 2, we used a sequence that was directly related to the force field itself, with a 6-element sequence of force field directions. While subjects were aware of the force field, the slight differences in force field direction were less obvious. Nonetheless, robust sequence learning also occurred in experiment 2. These findings indicate our brain can process sequence learning during sensorimotor learning tasks, even when the sequenced elements were subtle and closely integrated with the skill learning. In experiment 1, the anticipatory IEMG before the 12 reaches partially distinguished the sequence group from the random group but might not be a good indicator of sequence learning in this experiment paradigm. Because different muscle(s) showed anticipatory activity on different reach directions, it is difficult to evaluate sequence learning at the trial level this way. In contrast, reaction time can be expected to reflect sequence learning independent of reach direction.

### 4.2 Sensorimotor adaptation occurred under complex force fields

Participants were able to adapt to the force fields in both experiments in the current study. We used a distance-dependent force field to mimic the classic velocity-dependent force field characteristics (Shadmehr & Mussa-Ivaldi, 1994). Regardless of participants’ reaching speed, a distance-dependent force field provides the same perturbation experience to each individual on a given reaching path. This was important in the present study because the anticipatory information obtained from sequence learning could alter movement speed (Moisello et al., 2009). If the reaching speed would be quite different due to anticipation, then the perturbation applied by a velocity-dependent force field would differ across trials and participants as a result. Donchin and Shadmehr (2002) found participants were not able to learn random peak velocity force fields, another study also supported their findings (Mawase & Karniel, 2012). Therefore, using a distance-dependent force field, we ensured that all participants would experience the same peak sensorimotor perturbation, regardless of the reaching speed and added anticipatory information. Adaptation to the distance-dependent field followed the pattern we would expect with a velocity-dependent field; both types of perturbation initially caused large reaching errors that were gradually reduced through trial-and-error practice, with evidence of a negative aftereffect when the force field was abruptly removed (Sexton, Liu, & Block, 2019). This suggests that a distance-dependent force field can be used to investigate force field adaptation, even with the interaction of sequence learning.

In the first experiment of the current study, there were 12 different reaching directions. Unlike traditional center-out reaching tasks where the starting position of each reach is the same, the start and end position of each reach varied among four different targets on the four corners of a square. This made the task less straightforward to learn as the force field was counterclockwise to the reaching direction. For example, when reaching upward the participant would be pushed to the right by the force field, and when reaching downward the participant would be pushed to the left. Despite the task complexity, we still observed significant adaptation and washout in both sequence with force and random with force groups, indicating sensorimotor learning occurs robustly under these conditions.

The individual reaching directions examined in experiment 1 showed different adaptation magnitudes. Many force field adaptation tasks investigate only one reaching direction (e.g., forward reaching) (Mawase & Karniel, 2012), or average reaching errors from all directions (e.g., center-out reaching) (Criscimagna-Hemminger, Bastian, & Shadmehr, 2010). In experiment 1, we covered a wide range of 2-dimensional reach directions. The adaptation and aftereffect curves looked as expected for vertical and diagonal reach directions but were more flat for the horizontal directions. This was the case for both sequence and random with force groups. This might due to biomechanical considerations, with the upper limb joints more stable and able to absorb external perturbations during lateral movement.

In the second experiment, the reaching task was simpler in the sense that participants were asked to only reach upward, and the robot would bring their hand back to start position. However, the sequence component was hidden in the force field as the different force directions. All the force directions were rightward with only slight variation in angle, which made it harder to adapt to a changing force field. Though it is a novel type of force field adaptation task (we still used the distance-dependent force field), participants were able to adapt, indicating the brain was able to learn the changing environment despite multiple force directions.

### 4.3 Sequence facilitated sensorimotor learning

When adding a sequence component to force field adaptation, sequence learning and sensorimotor learning both occurred in both experiments. Moreover, we found evidences of sequence learning benefitting adaptation and retention in experiment 2. This suggests that the presence of sequence learning may be more likely to benefit sensorimotor learning if the elements being sequenced are the sensorimotor perturbation itself. In contrast, when the sequenced elements were the target positions in experiment 1, the same force field was experienced no matter what target position was presented next, so this sequence could be interpreted as less closely tied to the sensorimotor perturbation.

Aftereffect is an indicator of learning and retention of the force field perturbation that is assessed during the washout block, when force perturbation has been removed. Larger errors in the opposite direction (negative aftereffect) suggest the brain has robustly stored the learned sensorimotor changes (Shadmehr & Mussa-Ivaldi, 1994). A catch reach during adaptation that randomly turn off the perturbation is another way to detect negative aftereffect (Stockinger, Focke, & Stein, 2014). Though we did not find differences in the catch reach or in the washout block between groups in Experiment 1, in Experiment 2 we found significant group effects in both washout blocks. Specifically, the sequence group’s negative aftereffect was larger and more persistent (better retained) than the random group.

Early adaptation to a force field is substantial and does not differ greatly among typically developing adults (Lamothe et al., 2014; Krakauer & Mazzoni, 2011). We observed similar early adaptation rates between the two groups in experiment 1 as well as the groups in the first adaptation block of experiment 2 when predictive information was not available. However, in the second adaptation block of experiment 2, the SDF group kept the same sequence from the first adaptation block and adapted faster to the external perturbation than the random group. This suggests that the sequence component played an important role in re-learning the sensorimotor perturbation.

In Experiment 1, when comparing the perpendicular deviation on a trial level (averaged every 12 elements), we did not observe group differences. When examining individual reaching directions, there were differences in task performance among some reach directions, indicating the sequence of target appearance might have an influence on some reach directions but not all. In contrast with experiment 1, the sequence component in experiment 2 clearly affected adaptation in the individual force directions. This indicates that this form of sequence component, if added to force field adaptation, might assist the adaptation.

### 4.4 Implications and future directions

The current study combined two processes that have frequently been studied in isolation. Importantly, we found that sequence learning did not reduce sensorimotor learning and was even facilitatory in certain conditions. This has implications for how motor skills should be practiced and taught. For example, whether training an athlete or rehabilitating a stroke survivor, it may be advantageous to encourage repetition of all the elements of the skill in the correct sequence, rather than excessive practice of one element at a time in order to master the sensorimotor aspect first. Further research with more complex sensorimotor demands and more types of sequence will be needed to determine any specific recommendations.

Given that sequence and sensorimotor learning are thought to depend on many of the same brain regions, we did consider the alternative that adding sequence learning demands might be detrimental to sensorimotor learning. Specifically, we might have expected that adding a sequence element would exhaust neural resources available for sensorimotor learning, leading to reduced adaptation or aftereffect. However, literature directly comparing the two processes is quite limited, and meta analysis of separate neuroimaging studies of the two processes found only that some of the same brain regions such as SMA and premotor cortex are active in both types of study (Hardwick et al., 2013). The present results are inconsistent with the idea that sensorimotor and sequence learning share these neural substrates in any mutually exclusive fashion. The shared brain regions might not serve both processes simultaneously, but instead, gradually shift emphasis from one process to another, or each process might rely on separate populations of neurons within the same brain area.

We did not inform participants about the presence of a sequence. However, many of them noticed some amount of the sequence by the end of the experiment, especially when the sequence was target positions in experiment 1. This shift of implicit learning to explicit learning is common in sequence learning studies (Nissen & Bullemer, 1987). Although researchers tried to separate explicit and implicit learning in laboratory settings, we do not learn motor behaviors purely relying on only one of them in most of our real-life settings. In the present study, participants perceived the task to be purely a force field reaching task at the beginning. During this early stage of learning, motor areas were highly engaged in learning and understanding the force field (Richardson et al., 2006). As the participants gradually perceived the existence of a sequence to some degree, their reaching skills were already quite improved. At this stage, the brain areas such as the SMA and pre-SMA became more involved in the sequence learning aspect of the task (Roland et al., 1980). And when both processes were skilled, less neural resources might be required to connect the two processes; indeed, having shared brain regions might benefit more. As we can see in the second adaptation block in experiment 2, the re-learning of the force field was more efficient when having the predictive information from the previous adaptation block, and retention of force field learning was enhanced after the force field was turned off. This kind of interaction between explicit and implicit learning is supported by the relationship between involved brain areas (Ashe, Lungu, Basford, & Lu, 2006). Supplementary motor area (SMA) and pre-SMA became important during this time. This change of brain area engagement is similar in sensorimotor learning tasks (Hardwick et al., 2013).

The finding of the current study supports the way of learning in real life scenarios, that learning sequence and sensorimotor aspect together would not impair our learning. However, unlike research settings, in which explicit learning and implicit learning are more strictly separated, people rely on both to learn a sequence in real life. Future research should therefore investigate the effect of explicit sequence learning on sensorimotor adaptation. It is possible that explicit sequence learning could exhaust the sensorimotor resources, especially early in the task when sensorimotor learning is also more explicit. When both processes kick in, one might intentionally look for a sequence while trying to adapt to a new environment.

Although we did not find differences in sensorimotor adaptation with sequenced target positions, we cannot rule out target position as a sequence element that might affect sensorimotor learning. The sequence target positions we used in experiment 1 was relatively complicated when taking counterclockwise force field and the 12-element long sequence into account. We might observe differences if we make the targets less distinct from each other, for example, with only slight difference in reach angles. Similarly, based on experiment 2, we cannot conclude that force direction is the only element whose sequencing could facilitate sensorimotor learning. Force magnitudes could be another such variable, as it is related to the sensorimotor perturbation itself. Previous studies found participants were not able to adapt to randomly changing force magnitudes (Donchin & Shadmehr, 2002; Mawase & Karniel, 2012), so there may be limitations to this. A systematic study of different combinations of the two is needed.

A limitation of the study is lack of “incidental learning”. Many sequence learning tasks implement random catch trials among sequence blocks as a way to study sequence learning (Willingham et al., 1989; Wilkinson & Shanks, 2004). The reaction time or response time, if using a button-press paradigm, would gradually decrease after a few sequence blocks, but a random block would suddenly increase the reaction time or response time. When the previous sequence is resumed, the time would decrease again. This sudden change of time is an indicator of sequence learning. In the current study, we did not add such a random trial into the force field adaptation task, which limited us to comparing reaction time early and late in the session to evaluate sequence learning. However, we chose not to add such random trials due to the task complexity. If random trials were added into adaptation blocks, reaching error would likely increase for the sequence groups, making it difficult to compare adaptation between groups. Moreover, the number of reaches in the adaptation blocks would have had to increase to ensure the same opportunity for sequence learning as the random group, but then the sequence group would have longer force field exposure. In other words, we chose not to add a random trial to avoid unnecessarily complicating the interpretation. But with the findings from the current study that sequence learning did not impair force field adaptation, future studies could add random trials to an easier sensorimotor task to better investigate the role of sequence learning on sensorimotor learning.

## 5. Conclusion

The current study investigated the effect of adding sequence learning to a sensorimotor learning task in two different sequence paradigms. When the sequence was target positions in space, sequence learning did not influence the force field adaptation. However, when the sequence was force directions, adding the predictive information improved both retention and the rate of re-learning. Our findings indicate these two processes, when learnt together, are unlikely to monopolize each other’s neural resources, and might in turn facilitate each other in some conditions.

